# Inborn cardiorespiratory fitness and exercise training modulate brown adipose tissue function and plasticity in early life

**DOI:** 10.64898/2026.04.01.715665

**Authors:** Meagan S. Kingren, Daniel G. Sadler, Mary Barre, Lillie D. Treas, James D. Sikes, Steven L. Britton, Lauren G. Koch, Elisabet Børsheim, Craig Porter

## Abstract

This study aimed to determine the impact of inborn metabolic fitness and early life exercise training on whole body and brown adipose tissue (BAT) energetics. We carried out comprehensive metabolic phenotyping on 4-week old rats bred for high (high-capacity runner, HCR) and low (low-capacity runner, LCR) running capacity following randomization to voluntary wheel running (VWR) or control (CRTL) for 6-weeks. High-resolution respirometry and untargeted proteomics were then employed to determine the impact of inborn fitness and early life exercise on BAT function. When accounting for differences in body mass, early life exercise (VWR) resulted in greater basal and total energy expenditure, irrespective of strain (*P* < 0.0001 for both). Both leak and uncoupling protein 1 (UCP1) dependent respiratory capacities in isolated BAT mitochondria were greater in rats randomized to VWR compared to CTRL in both HCR (*P* < 0.01) and LCR (*P* < 0.05) strains. Similarly, mitochondrial sensitivity to the UCP1 inhibitor GDP was greater in both HCR (*P* < 0.01) and LCR (*P* < 0.05) rats randomized to VWR versus control. The BAT proteome differed in CTRL HCR and LCR rats, were there was enrichment in proteins related to branched chain oxidation and mitochondrial fatty acid oxidation in HCR rats. VWR remodeled the BAT proteome, where 151 proteins were differentially expressed in LCR BAT and 209 differentially expressed in LCR BAT following VWR. In both stains, there was an enrichment in proteins related to metabolism mitochondrial function in response to VWR. However, when comparing strains, 39 proteins were differentially expressed in BAT in HCR rats compared to LCR rats in response to VWR. These proteins were related to carboxylic acid and amino acid metabolism. Collectively, inborn fitness impacts body mass and composition, exercise behaviors, and the BAT proteome in early life. Early life exercise alters whole body and BAT energetics irrespective of inborn fitness, augmenting basal and total energy expenditure and BAT thermogenic capacity and function.

## Introduction

A vast majority of children (81%) worldwide fail to meet WHO physical activity guidelines, and ∼31% of adults are exercise insufficient^1,2^. The rate of exercise insufficiency has increased significantly over the last two decades^3^ and correlates with increased obesity and metabolic disease prevalence^4–6^. Further, the age of obesity onset has shifted to younger ages, and in the United States, approximately 1 in 3 children are overweight or obese^7^. Childhood obesity is associated with poor lifelong health outcomes such as cardiovascular disease^8,9^, delayed cognitive development^10^, and type 2 diabetes mellitus^11,12^. Childhood obesity confers a five-fold greater risk of obesity in adulthood, and nearly 70% of obese adolescents remain obese over the age of 30^13^. This underscores the need for strategies to mitigate childhood obesity and metabolic disease risk.

Greater aerobic fitness is associated with reduced risk for hypertension, obesity, diabetes, stroke, and cancer^14–16^. Indeed, high cardiorespiratory fitness in adults is associated with a delayed onset of common chronic diseases^17^, and early-life exercise is associated with lower rates of body fat acquisition in early adolescence^18^. Further emphasizing the protective effects of exercise in animal models, early-life exercise extends the health span of mice that were otherwise sedentary^19^. While aerobic exercise can improve measures of cardiorespiratory fitness (CRF), up to 60% of CRF is attributable to genetic factors^20,21^. To evaluate this inborn component of cardiorespiratory fitness, two-way selective breeding of rats based on maximal treadmill running performance generated high- and low-capacity runner (HCR/LCR) offspring with roughly an 8-fold difference in exercise capacity^22^. HCR rats exhibit better metabolic function and protection against cardiometabolic diseases^23–25^. Our recent work found inborn fitness levels influence molecular adaptations to early life exercise in skeletal muscle and liver, where early-life exercise only partially rescues the effects of low inborn fitness^26^.

Brown adipose tissue (BAT) has recently re-emerged as a tissue of interest with regards to mammalian energetics and metabolic health. Through uncoupling protein-1 (UCP1), BAT is able to dissipate mitochondrial membrane potential as heat, rather than using it for ATP synthesis^27^. As such, BAT UCP1 modulates non-shivering thermogenesis—and thus energy expenditure—in mammals, including humans. Indeed, BAT activation via cold exposure can acutely increase energy expenditure and alter glucose and lipid metabolism in humans. Like cold exposure, exercise activates the sympathetic nervous system, stimulating the release of norepinephrine, activating BAT. Further, in humans, lower BMI is associated with greater BAT volume^28^. However, whether BAT function and its responsiveness to exercise is influenced by parental cardiorespiratory function remains unknown.

Here we sought to evaluate the following hypothesis: BAT bioenergetics and BAT remodeling in response to early life exercise training are influenced by parental cardiorespiratory fitness, where BAT from offspring born to parents with higher cardiorespiratory fitness have greater thermogenic capacity and exhibits greater plasticity in response to exercise training. To test this hypothesis, we utilized rats selectively bred for low and high capacity running and randomized these juvenile rats to six weeks of voluntary wheel running (VWR) or control (CTRL).

## Methods

### Regulatory Approval

All animal procedures were approved by the University of Arkansas for Medical Sciences Institutional Animal Care and Use Committee (protocol no. 4167). All animal experiments were carried out in line with the ARRIVE guidelines for animal research.

### Animals

An *a priori* power analysis was carried out as described previously where we determined with an actual power of 0.82 that nine animals per group would be sufficient to detect intragroup differences in blood glucose levels. We utilized n=10 per group (50% male) for this study. Rats bred for high and low running capacity (HCR/LCR) were obtained from the University of Toledo (OH, USA) following rapid quarantine at Charles River Laboratories (Wilmington, MA, USA) before arrival at our facility. Upon arrival at ∼16-20 weeks of age, breeding pairs from generation 47 were housed at 24°C on a standard 12/12h light/dark cycle (7:00 am lights on) with *ad libitum* access to food (ProLab RMH 1800; 21.1% protein, 13.8% fat, and 65.1% carbohydrate; 4.1 kcal/g; LabDiet, St. Louis, MO, USA) and drinking water. Strains were housed separately. Virgin male and female rats were paired for breeding, and upon confirmation of pregnancy, males were removed from the cages. Breeding pairs produced a total of 16 litters that were subsequently weaned at 26 days of age. Following weaning, two or three offspring were randomly selected based on body mass to provide a total of 20 rats per strain (50% male), and the remaining pups were humanely euthanized. Pups selected for the study (n=20 per strain, 50% male) were randomized to control (CTRL) or voluntary wheel running (VWR) groups based on body mass to create final groups of n=10 per strain per condition (50% male). Following group allocation, rats were singly housed with or without access to running wheels in standard home cages (Techniplast Activity Systems; Starr Life Sciences Corp., Oakmont, PA, USA) for 6 weeks during which VWR was monitored in real time using VitalView activity software (Starr Life Sciences Corp.). Following conclusion of the VWR period, running wheels were locked and rats were euthanized within two hours using a rising concentration of CO_2_.

### Tissue Collection

Rats were euthanized upon conclusion of the six-week VWR period. The entire intrascapular BAT pad was collected and placed in ice-cold sucrose isolation buffer (225 mM mannitol, 75 mM sucrose, and 0.2 mM EDTA) for subsequent mitochondrial isolation.

### Mitochondrial Isolation

Mitochondrial isolation from BAT was based on our protocol described previously^26^. Briefly, the entire BAT pad was minced prior to homogenization with a glass polytetrafluoroethylene pestle for five strokes in ice-cold sucrose isolation buffer (25 mM mannitol, 75 mM sucrose, and 0.2 mM EDTA). The resulting homogenate was filtered through one layer of cheesecloth and centrifuged at 8500 *g* for 10 min at 4°C. Following discard of the hard-packed fat layer and supernatant, the pellet was resuspended in a volume of 10 mL of sucrose buffer and centrifuged at 800 *g* for 10 min at 4°C. The resulting supernatant was centrifuged at 8500 *g* for 10 min at 4°C before resuspension of the subsequently formed mitochondrial pellet in 0.3% fatty-acid-free bovine serum albumin supplemented sucrose buffer and centrifugation at 8500 *g* for 10 min at 4°C. Following resuspension in 5 mL of KCl-TES solution and centrifugation at 8500 *g* for 10 min at 4°C, the osmotically-sound, fatty-acid-free mitochondrial pellet was resuspended for later high-resolution respirometry analysis and storage for quantitative proteomics.

### High Resolution Respirometry

A final concentration of 0.05mg/mL of BAT mitochondrial isolate were evaluated in MIR05 buffer (0.5mM EGTA; 3 mM MgCl_2_; 0.5 M K-lactobionate; 20 mM taurine; 10 mM KH_2_PO_4_; 20 mM HEPES; 110 mM sucrose; 1 mg/ml essential fatty acid free bovine serum albumin) maintained at 37°C for respiration analysis in a 2 mL Oxygraph O2K Respirometer chamber (Oroboros Instruments, Innsbruck, Austria). Leak respiration was determined following the addition of glycerol-3-3phosphate (10mM). Thereafter, GDP was titrated at the following concentrations to inhibit UCP1 (final chamber concentration in (brackets): 10µM (10µM), 10µM (20µM), 20µM (40µM), 40µM (80µM), 80µM (160µM), 160µM (320µM), 320µM (640µM), 640µM (1280µM), 1280µM (2560µM), 2560µM (5120µM). Oxygen consumption rate per second was calculated per 100 μg of mitochondrial protein following the addition of saturating concentrations and was recorded at 2-4s intervals (DatLab, Oroboros Instruments, Innsbruck, Austria).

### Quantitative Proteomics

Using the IDeA National Resource for Quantitative Proteomics we carried out proteomics analysis as previously described^29^. Briefly, total protein from tissue (*n*=10/group/tissue) was concentration-matched, reduced, alkylated, and purified by chloroform/methanol extraction prior to digestion with sequencing grade modified porcine trypsin (Promega, Madison, WI, USA). Tryptic peptides were then separated by reverse phase XSelect CSH C18 2.5 um resin (Waters, Milford, MA) on a column using an UltiMate 3000 RSLCnano system (Thermo). Using a 60 min gradient from 98:2 to 65:35 buffer A:B ratio (Buffer A = 0.1% formic acid, 0.5% acetonitrile; Buffer B = 0.1% formic acid, 99.9% acetonitrile), peptides were eluted and then ionized by electrospray (2.4kV) and mass spectrometric analysis was carried out on an Orbitrap Exploris 480 mass spectrometer (Thermo). To assemble a chromatogram library, six gas-phase fractions were acquired (Orbitrap Exploris with 4 m/z DIA spectra, 4 m/z precursor isolation windows at 30,000 resolution, normalized AGC target 100%, maximum inject time 66 ms) using a staggered window pattern from narrow mass ranges using optimized window placements. Precursor spectra were acquired after each DIA duty cycle, spanning the m/z range of the gas-phase fraction (i.e. 496-602 m/z, 60,000 resolution, normalized AGC target 100%, maximum injection time 50 ms). For wide-window acquisitions, the Orbitrap Exploris was configured to acquire a precursor scan (385-1015 m/z, 60,000 resolution, normalized AGC target 100%, maximum injection time 50 ms) followed by 50 x 12 m/z DIA spectra (12 m/z precursor isolation windows at 15,000 resolution, normalized AGC target 100%, maximum injection time 33 ms) using a staggered window pattern with optimized window placements. Precursor spectra were acquired after each DIA duty cycle.

### Proteomics Data Analysis

To obtain a comprehensive proteomic profile, obtained data were searched using an empirically corrected library and quantitative analysis performed. Proteins were identified and quantified using EncyclopeDIA and visualized with Scaffold DIA using 1% false discovery thresholds at both the protein and peptide level ^30^. Protein exclusive intensity values were assessed for quality and normalized using ProteiNorm ^31^. The data was normalized using Cyclic Loess and statistical analysis was performed using Linear Models for Microarray Data (limma) with empirical Bayes (eBayes) smoothing to the standard errors ^32^. Only proteins expressed in at least 2/3 of samples were considered in analysis. Differentially abundant proteins (DAPs) were those with an *adj. P*-value ≤0.05 and an absolute fold change ≥1.5. UniProt IDs (protein) were then converted to Ensembl ID (mouse gene) for enrichment analysis using gProfiler^33^. Differnetial analyses were carried out in R 4.5.2+ using ProteoDA^34^.

### Statistical information

Data are presented as mean ± standard deviation. Two-way analyses of variance (ANOVA) were performed to detect strain and exercise effects. Pairwise comparisons (for breeder data) were calculate using Šidák’s multiple comparisons test. Analyses were carried out in GraphPad Prism 10 (GraphPad software Inc., San Diego, CA, USA). Proteomics analyses were carried out in R as described above. For IC50 values of UCP1-driven respiration following GDP titration, we calculated dose response curves using a nonlinear regression assuming a four-parameter sigmoidal relationship to log[GDP] with a fixed bottom respiration value, similar to prior work by Musiol and colleagues^35^. To compare the models, an extra sum-of-squares F test for the LogIC50 was performed. Single replicate curves were calculated as well.

## Author Contributions

C.P., E.B., S.B., and L.K. conceived the study and designed the experiments; D.G.S., M.B., L.D.T., T.R., J.D.S., and C.P. carried out experimentation; D.G.S. carried out analysis on metabolic phenotyping data. M.S.K. analyzed and interpreted the rest of the data and wrote the manuscript. All authors reviewed, edited, and approved the final version of the manuscript.

## Results

### Inborn fitness levels dictate exercise physiology in rats

Rats bred at the University of Toledo for high and low running capacity (HCR (n=16, 50% male) and LCR (n=16, 50% male)) were used to generate HCR and LCR offspring for this current study. As previously reported, HCR breeders ∼5-fold greater time to exhaustion (67.0 ± 5.0 vs 12.5 ± 1.3 mins; *P* < 0.001), ∼11-fold greater total distance (1764 ± 220.2 vs 157.6 ± 20.7 m; *P* < 0.001) and ∼2.5-fold greater running speed (39.8 ± 3.0 vs 15.2 ± 1.6; *P* < 0.001) compared to LCR breeders^26^. Upon arrival to our facility at 16-20 weeks of age (n=16 per strain; 50% male), all HCR and LCR breeders were phenotyped, where both male and female HCR breeders were smaller (275.1 ± 56.5 vs 359.9 ± 117.9 g body mass; *P* = 0.0144), leaner (232.2 ± 40.7 vs 298.0 ± 82.8 g lean mass; *P* = 0.0087) and had greater glucose control in response to a glucose tolerance test (strain main effect *P* < 0.001; lower area-under-the-curve, *P* = 0.0246)^26^. A total of eight virgin females from both HCR and LCR strains were paired with males from the same strain at approximately 24 weeks of age (a different sire was used in each breeding pair). Sires were removed from the cage once pregnancy was confirmed. Upon delivery of pups, litters were housed with their mothers until postnatal day 26. Thereafter, pups were randomly selected from all litters (no more than n=2 from each litter) and randomized within each strain to either voluntary wheel running (VWR) or control (CTRL) (n=10 per group; 50% male).

### Early-life exercise partially rescues indices of low inborn fitness

Per our previous report^26^ HCR rats had lower body mass, absolute fat mass, and relative fat mass compared to LCR rats (**Supplemental Figure 1**). Further, strain and exercise effects emerged upon examination of changes in body composition over the six-week VWR period (Figure 1). HCR gained less mass, regardless of exercise status (120.1 ± 22.1 vs 182.2 ± 24.7 g body mass; strain main effect *P* < 0.001) compared to LCR, while VWR served to further limit body mass accretion in HCR (134.6 ± 42.5 vs 167.7 ± 45.2 g body mass; VWR main effect *P* = 0.02) **Figure 1A**). HCR gained less fat and lean mass compared to LCR irrespective of exercise. There was an interaction between greater inborn fitness and VWR where there was less fat mass deposition in HCR rats that exercised (*P* = 0.03 interaction effect and strain main effect, *P* < 0.0001 for VWR main effect; **Figure 1B**). Within sedentary rats, HCR-CTRL gained less fat mass than LCR-CTRL (9.9 ± 5.4 vs 17.9 ± 8.3 g; *P* = 0.02), indicating higher inborn fitness protects against early-life fat deposition. Exercise had a significant effect in LCR in terms of adipose tissue accretion, where LCR-VWR had over 4-fold less fat mass gain compared to LCR-CTRL (4.2 ± 4.5 vs 17.9 ± 8.3 g fat mass; *P* < 0.001). While fat mass accretion was numerically lower HCR-VWR vs HCR-CTRL, this was not statistically significant. Differences in lean mass deposition further explain the nearly 30% lower body mass in HCR compared to LCR (**Figure 1C**). While VWR did not have a main effect on lean mass deposition, strain did. LCR-CTRL had a greater change in lean mass than HCR-CTRL (157.4 ± 10.4 vs 104.2 ± 18.1 g lean mass; *P* < 0.001). VWR had a strain-dependent impact on glucose tolerance, as we previously published^26^. VWR resulted in lower blood glucose levels in LCR, but not HCR^26^. Strain effects were also seen in voluntary wheel running distance, where HCR-VWR ran more than LCR-VWR beginning as early as the second week of exercise training (*P* < 0.0001; **Figure 1D-E**). Upon completion of the early-life exercise period, HCR-VWR averaged nearly 3.5-fold more m/day than LCR-VWR (8375.5 ± 5478.5 vs 2395.5 ± 1293.8 m/day; *P* = 0.0002). Emphasizing that HCR are predisposed to be more active, HCR-CTRL ambulated about their home cage more than LCR-CTRL (156.2 ± 42 vs 49.7 ± 23.8 m; *P* < 0.001; **Figure 1F**), spending approximately twice as much time ambulating around the cage (18.8 ± 4.3 vs 8.3 ± 3.5% walking time; *P* < 0.001; **Figure 1G**). Despite LCR rats being significantly larger than HCR rats, there were no differences in total energy expenditure, basal energy expenditure, or peak active energy expenditure (**Supplemental Figure 1**). To better understand the relationship between body mass and energy expenditure, we carried out an analysis of covariance (ANCOVA). When accounting for body mass and strain, HCR rats had greater TEE than LCR, despite LCR being ∼30% larger (*P* = 0.0055; **Supplemental Figure 1**). Indeed, when accounting for body mass and group, a significant effect of early-life exercise on TEE emerged (**Figure 1H**). HCR-VWR had the greatest TEE, followed by LCR-VWR, HCR-CTRL, and LCR-CTRL (ANCOVA *P* < 0.0001). We then examined BEE, determined as the lowest 30 mins or energy expenditure during the light cycle. We found that VWR led to higher BEE (ANVOCA *P* < 0.0001; **Figure 1I**). However, when accounting for BEE by body mass and strain without exercise, strain alone did not influence BEE (**Supplemental Figure 1**), further indicating that exercise training is critical in BEE in HCR and LCR rats. Together, these data indicate early-life exercise results in greater energy expenditure, where greater inborn confers increased exercise capacity to drive up TEE. When examining BEE as a proportion of TEE, there is a significant effect of VWR, where the non-BEE proportion of energy expenditure was greater in VWR compared to CTRL (**Figure 1J**).

**Figure 1.**
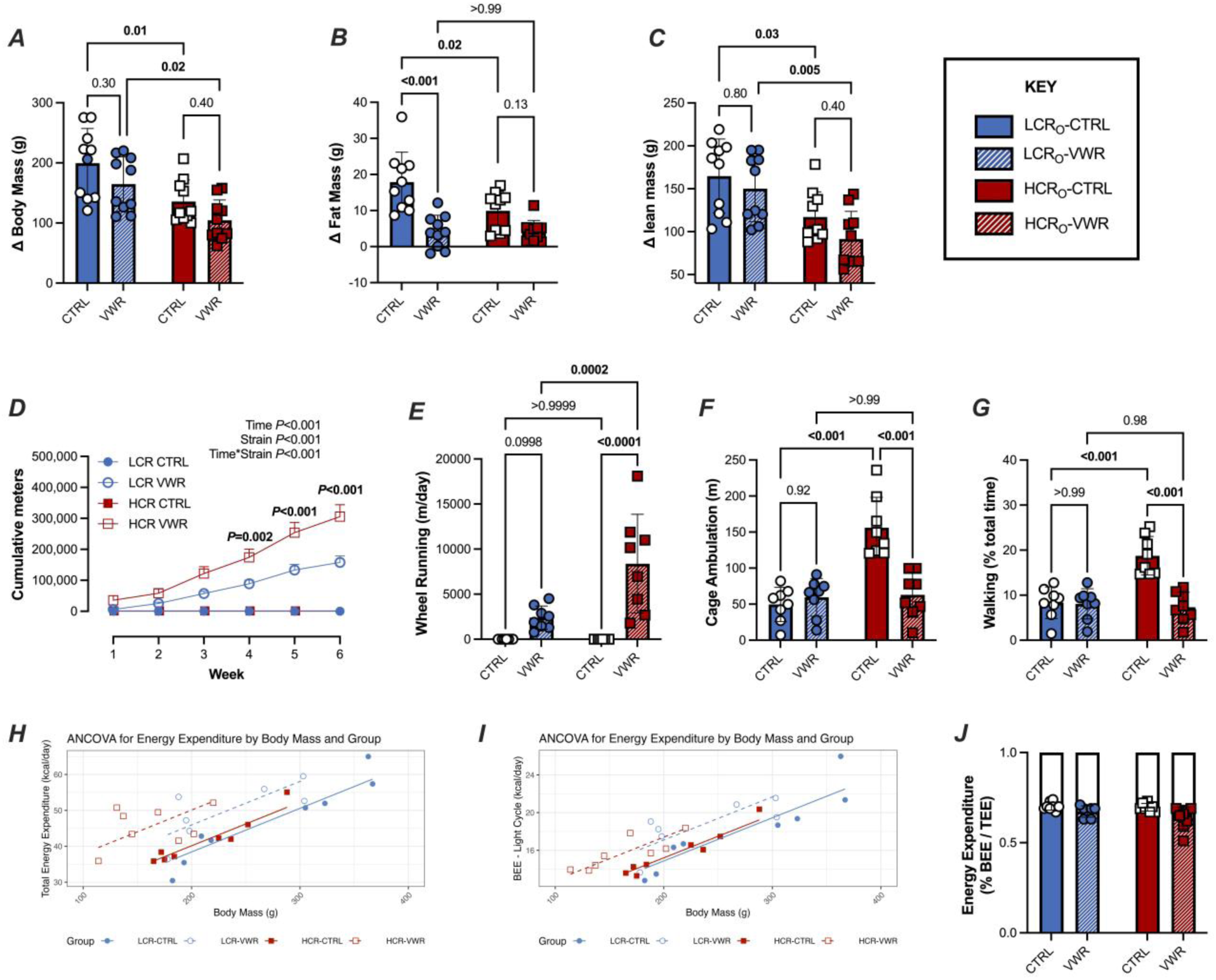
Exercise partially rescues measures of poor metabolic health imparted by low inborn fitness. Echo-MRIs were carried out prior to VWR treatment and six-weeks later before euthanasia to determine differences in body composition over time. Rates of (**A**) body mass, (**B**) fat mass, and (**C**) lean mass deposition are shown. (**D-E**) Voluntary wheel running activity was monitored, as was (**F**) ambulation about the home cage and (**G**) time spent walking about the home cage. Energy expenditure was also monitored using specialized cages to quantify TEE and BEE. An ANCOVA to evaluate the effect of body mass and strain on (**H**) TEE and **(I**) BEE in the light cycle are shown. (**J**) BEE as a proportion of TEE is shown. Abbreviations: TEE—total energy expenditure; BEE—basal energy expenditure. Differences quantified using two-way ANOVA with exceptions of (D), where a three-way ANOVA was used to identify the effects of strain and exercise over time and (H-I), where ANCOVAs were used to identify the effects of body mass and group on energy expenditure.

### Exercise training alters inborn fitness-driven BAT mitochondrial function

VWR led to lower BAT depot mass in LCR (233.9 ± 39.1 vs 276.5 ± 50.1 g; *P* = 0.0323) and HCR (202.0 ± 32.9 vs 286.0 ± 47.0 g; *P* < 0.0001) compared to CTRL (exercise main effect *P* < 0.0001; **Figure 2A**). When normalized to total body mass, there was a significant strain effect on BAT pad size, where HCR had larger BAT pads than LCR (0.134 ± 0.01 vs 0.098 ± 0.01 mg/g body mass; *P* = 0.0027; **Figure 2B**), HCR-CTRL had less mitochondrial protein compared to LCR-CTRL (6.0 ± 2.5 vs 9.6 ± 2.7 μg mitochondrial protein per mg homogenized BAT; *P* = 0.0221; **Figure 2C**). HCR-VWR saw elevated mitochondrial protein concentration compared to HCR-CTRL (9.3 ± 3.5 vs 6.0 ± 2.5 μg; *P* = 0.0308), though no changes were seen as a result of VWR in LCR (**Figure 2C**). There was a strain*exercise interaction effect on mitochondrial protein concentration (*P* = 0.0192). Once normalized to BAT tissue mass, total BAT depot mitochondrial protein levels were greater in LCR-CTRL versus HCR-CTRL (2582.9 ± 656.6 vs 1770.6 ± 959.2 μg; *P* = 0.0279; **Figure 2D**), though no broader strain main effect was detected. While no main effect of exercise was seen, LCR-VWR had a lower total depot protein level than LCR-CTRL (1756.7 ± 809.1 vs 2582.9 ± 656.6 μg; *P* = 0.0255).

**Figure 2.**
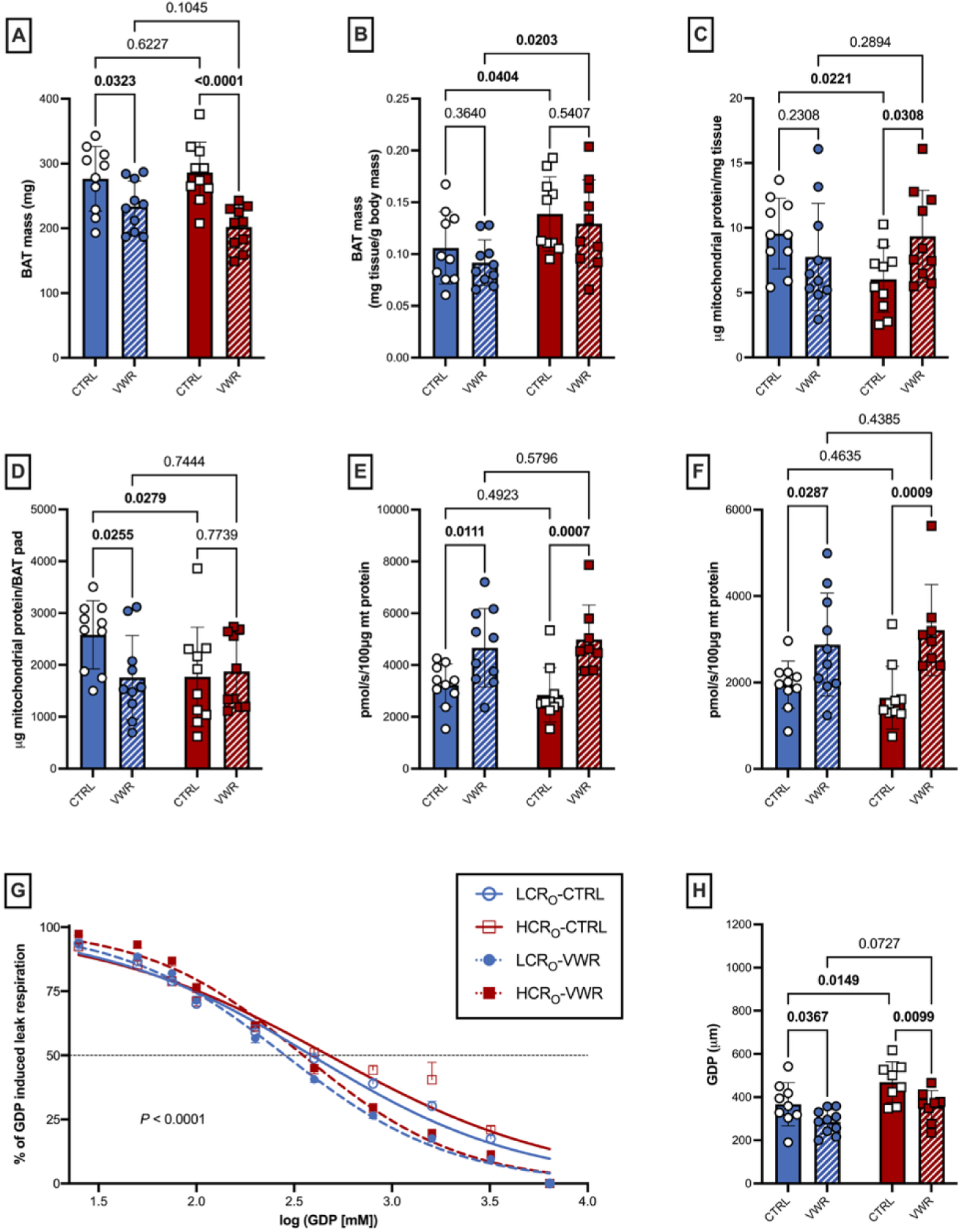
Exercise emphasizes inborn fitness related mitochondrial function. (**A**) BAT depot mass, (**B**) BAT mass normalized to total body mass (**C**) mitochondrial protein per mg of homogenized tissue, (**D**) mitochondrial protein normalized to BAT mass to obtain total depot protein concentration, (**E**) mitochondrial leak respiration, (**F**) UCP1-driven respiration, (**G-H**) BAT IC50 for HCR and LCR subjected to VWR or CTRL for 6 weeks. All statistics are presented as mean ± standard deviation. Comparisons to evaluate strain, exercise, and strain*exercise interaction effects were determined by 2-way ANOVA except for grouped IC50 curves (g). IC50 curves were determined using nonlinear regression of the log transformed GDP concentration values.

Using high-resolution respirometry (HRR; Oxygraph O2k, Oroboros Instruments), we examined mitochondrial function in BAT mitochondrial isolates. Through HRR, we quantified leak respiration supported by glycerol-3-phosphate and found a VWR main effect (*P* < 0.0001; **Figure 2E**). No strain effect was identified. Leak respiration was ∼1.4-fold greater in LCR-VWR (3222.9 ± 823.0 vs 4667.4 ± 1516.9 pmol/s/100μg mt protein; *P* = 0.0111) and ∼1.7-fold greater in HCR-VWR (2849.7 ± 1049.1 vs 4986.4 ± 1333.0 pmol/s/100μg mt protein; *P* = 0.0007) compared to their respective controls. We then examined UCP1-dependent respiration, where a significant VWR main effect was detected (*P* = 0.0002; **Figure 2F**). UCP1-dependent respiration was elevated ∼1.5-fold in LCR-VWR (1949.1± 547.6 vs 2876.4 ± 1191.9 pmol/s/100μg mt protein; *P* = 0.0287) and ∼1.9-fold in HCR-VWR (3213.8 ± 1052.4 vs 1648.2 ± 757.5 pmol/s/100μg mt protein; *P* = 0.0009) versus CTRL.

To further investigate the differences in UCP1 due to the combined effects of inborn fitness and early-life exercise, we compared changes in respirations following titrations of the UCP1 inhibitor GDP. Milli-Molar concentrations are typically employed to fully inhibit UCP1 in isolated BAT mitochondria. Here, we employed GDP titrations beginning in the µM range to represent a physiological intracellular GDP concentration, escalating to a final concentration of ∼5mM to ensure a maximal response to GDP. This allowed us to assay BAT UCP1 sensitivity to GDP and calculate the IC50 of GDP for UCP1 in isolated BAT mitochondria. The IC50, or the concentration of GDP where UCP1 supported respiration was reduced by half, was evaluated using a nonlinear regression with an extra sum-of-squares F test to identify that the LogIC50 was different across groups (*P* < 0.0001; **Figure 2G**). Main effects for VWR (*P* = 0.0014) and strain (*P* = 0.0037) were detected when comparing individual IC50 replicate curves (**Figure 2H**). Exercise resulted in lower IC50s for both LCR-VWR (285.4 ± 55.6 vs 367.3 ± 99.8 μM GDP; *P* = 0.0367) and HCR-VWR (357.3 ± 73.0 vs 469.5 ± 93.9 μM GDP; *P* = 0.0099) versus their respective CTRL group. HCR-CTRL had a greater IC50 (469.5 ± 93.9 vs 367.3 ± 99.8 μM GDP; *P* = 0.0149). These results indicate inborn fitness and early life exercise alter UCP1 function, specifically changing its sensitivity to inhibitory purine nucleotides.

### Inborn fitness levels dictate protein expression

To describe the proteomic responses to VWR in the context of high and low levels of inborn fitness, we utilized untargeted semi-quantitative proteomics in mitochondria isolated from brown adipose tissue (BAT). A total of 2,592 proteins were detected with a false discovery rate (FDR; *q*) less than 1% at the protein and peptide levels. After filtering proteins expressed in at least 66% of biological replicates, 2,500 proteins remained for subsequent analyses. Intensity data was normalized using cyclic loess (**Supplemental Figure 2**), and statistical analysis was performed using linear models for microarray data (limma) with empirical Bayes (eBayes) smoothing to the standard errors to determine differential proteins resulting from inborn fitness. Proteins with a Benjamini-Hochberg false discovery adjusted *p* ≤ 0.05 and an absolute fold change (FC) ≥ 1.5 were considered differentially abundant (DAPs).

We first determined the effects of inborn fitness in HCR-CTRL and LCR-CTRL groups, where rats lacked access to running wheels and could only ambulate about their home cage. We identified 27 DAPs between strains, where 12 proteins were more abundant and 15 proteins were less abundant in HCR-CTRL compared to LCR-CTRL (**Figure 3A**). Enrichment of Ingenuity Pathway Analysis (IPA) Canonical Pathways were calculated to determine potentially activated or inhibited signaling and metabolic pathways. HCR-CTRL demonstrated enrichment in isoleucine and valine degradation, branched chain amino acid (BCAA) catabolism, mitochondrial fatty acid β-oxidation, and thyroid hormone biosynthesis relative to LCR-CTRL. KEGG and GO enrichment terms were similar, indicating valine, leucine, and isoleucine degradation, pantothenate and CoA biosynthesis, isoleucine catabolic processes, BCAA metabolism, and the mitochondrion terms were all more enriched in HCR-CTRL versus LCR-CTRL (**Figure 3B**).

**Figure 3.**
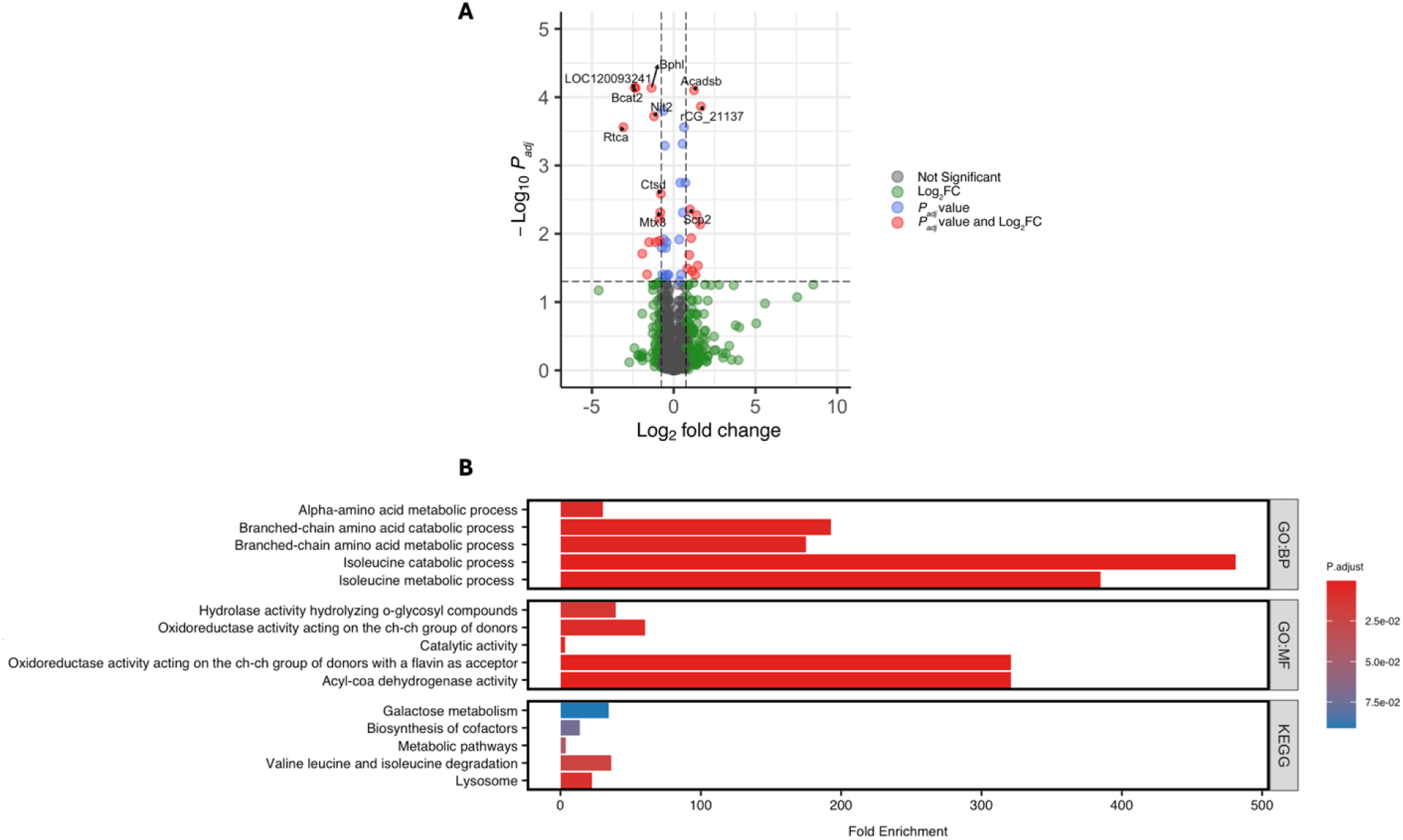
Inborn fitness alters the brown adipose tissue mitochondrial proteome. (**a**) Volcano plot showing differentially abundant proteins when comparing HCR-CTRL to LCR-CTRL. The top 10 DAPs are labeled. (**b**) DAPs were subjected to enrichment analyses, and the top 5 terms for GO:BP, GO:MF, and KEGG are shown. Abbreviations: DAP—Differentially abundant protein; GO:BP—Gene Ontology: Biological Processes, GO:MF—Gene Ontology: Molecular Function; KEGG—Kyoto Encyclopedia of Genes and Genomes.

### Early life exercise further alters the proteome

Early life exercise was modelled using 6 weeks of voluntary wheel running (VWR) in both HCR and LCR rats. VWR imparted notable proteome remodeling, regardless of inborn fitness level (**Figure 4**). In total, 151 DAPs were identified in LCR BAT in response to VWR. Of these DAPs, 68 were more abundant and 83 were less abundant in LCR-VWR compared to LCR-CTRL. IPA Canonical Pathway analysis indicated VWR results in altered mRNA processing and subsequent RNA post-transcriptional modifications, as well as impaired triglyceride metabolism, altered calcium handling, and decreased integrin to cytoskeleton signaling. GO and KEGG enrichment analyses corroborate these findings, with GO:BP and GO:CC terms indicating more abundant DAPs drive altered metabolic processes and less abundant DAPs drive altered mRNA and muscle system processes and enrichment of muscle-related cellular components.

**Figure 4.**
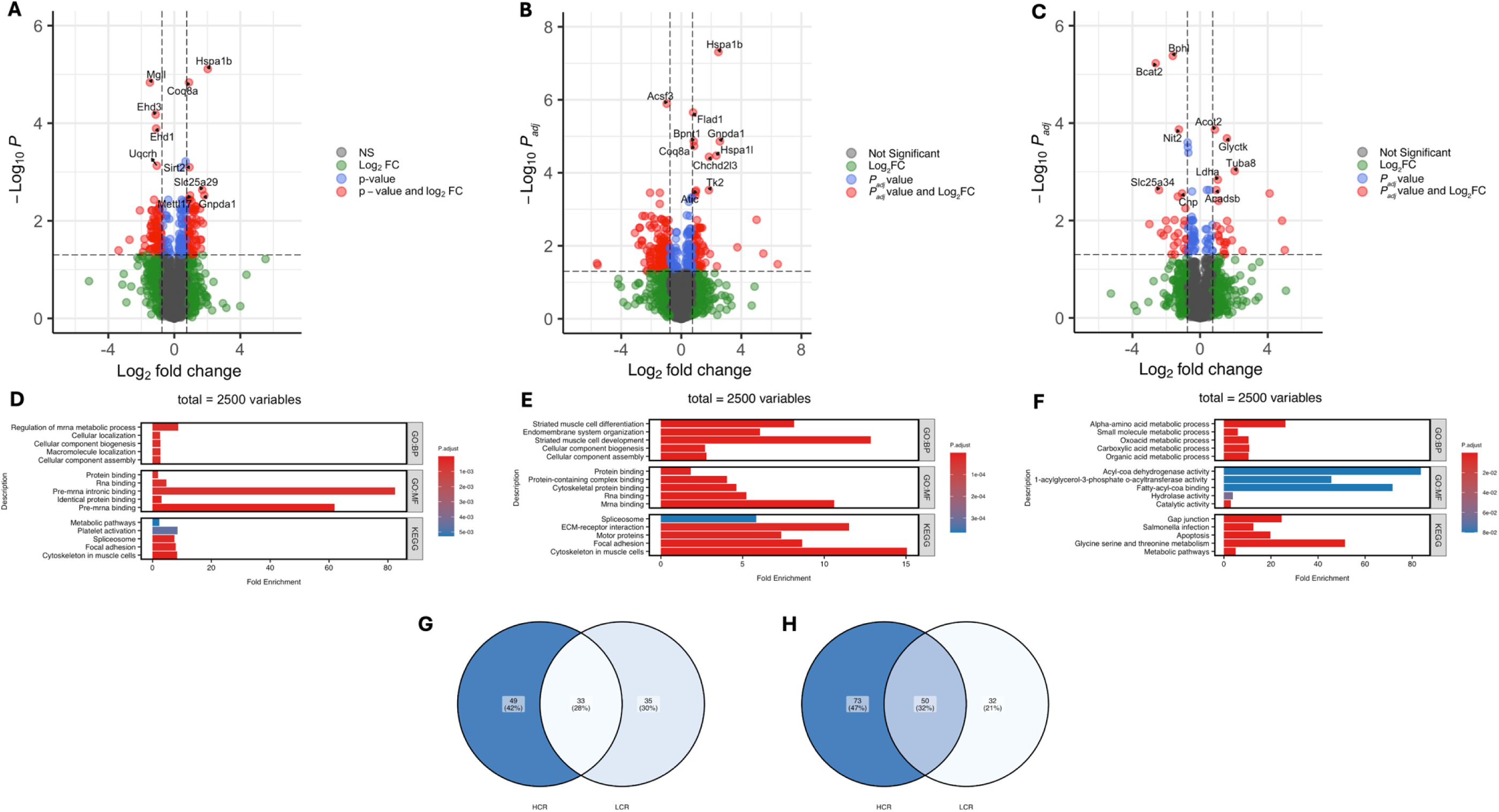
Exercise further alters the inborn fitness influenced BAT mitochondrial proteome. (**a-c**) Volcano plot showing DAPs for LCR-VWR versus LCR-CTRL, HCR-VWR versus HCR-CTRL, and HCR-VWR versus LCR-VWR are shown. The top 10 DAPs are labeled. (**d-f**) DAPs were subjected to enrichment analyses, and the top 5 terms for GO:BP, GO:MF, and KEGG are shown for LCR-VWR versus LCR-CTRL, HCR-VWR versus HCR-CTRL, and HCR-VWR versus LCR-VWR, respectively. Venn diagrams depicting the common DAPs upregulated (**g**) and downregulated (**h**) by exercise for HCR-VWR and LCR-VWR versus their respective CTRL groups are shown. Abbreviations: DAP—Differentially abundant protein; HCR—High capacity runner offspring rats; LCR—Low capacity runner offspring rats; VWR—Voluntary wheel running; CTRL—Sedentary control; GO:BP—Gene Ontology: Biological Processes, GO:MF—Gene Ontology: Molecular Function; KEGG—Kyoto Encyclopedia of Genes and Genomes.

The effect of VWR in HCR resulted in 209 DAPs, where 82 proteins were more abundant and 127 were less abundant in HCR-VWR compared to HCR-CTRL. The most significantly altered Canonical Pathways indicated VWR results in predicted decreases in BAT striated muscle contraction and muscle filament signaling, extracellular matrix organization, endocytosis signaling, and integrin cell surface interactions. More abundant DAPs are largely related to the mitochondrion and demonstrate enrichment in GO terms pertaining to small molecule metabolic processes, while less abundant DAPs appear to drive alterations in cytoskeletal protein binding, mRNA and muscle system processes, and muscle organization.

To further evaluate the influence of inborn fitness on the proteomic response to exercise, we compared HCR-VWR and LCR-VWR groups and found 49 DAPs (24 more abundant and 25 less abundant). Only 10 (15.2%) of these DAPs were commonly differential with the 27 DAPs found when comparing strains in CTRL rats. These 10 common DAPs related to BCAA catabolism and metabolic processes (GO:BP and KEGG). The 39 unique DAPs altered only in HCR-VWR pertained to carboxylic acid metabolism and caveola assembly (GO:BP), as well as glycine, serine, and threonine metabolism (KEGG) (**Figure 4**). These results indicate early-life exercise can magnify some proteomic differences resulting from inborn fitness.

## Discussion

The heterogeneous response to exercise is due largely to the altered responses of molecular transducers of physical activity^36,37^, including inborn fitness levels^29,38–41^. Building upon our previous study examining the effects of early-life exercise in LCR and HCR rats on the muscle and liver proteome and transcriptome^26^, our study herein profiled BAT in the context of inborn CRF and responses to early life exercise training. BAT is being increasingly examined as a potential therapeutic target for obesity and cardiometabolic disease due to its roles in non-shivering thermogenesis and as a metabolic sink for glucose, fatty acids and branched chain amino acids (BCAAs)^42–45^. BAT activation is associated with lower BCAA levels in both mice and humans^46^. Further, in humans, BAT is independently associated with healthier fat distribution, lower blood glucose levels, increased HDL, decreased hepatic steatosis, lower triglycerides, and diminished type 2 diabetes mellitus prevalence^47,48^. We therefore sought to examine the role of parental fitness in BAT physiology, as well as how early-life exercise can alter BAT function and proteome remodeling in rats born to parents with either high or low cardiorespiratory fitness. Our data suggest: (1) early life exercise exerts comparable changes in whole body energy expenditure and BAT bioenergetics, and (2) higher inborn fitness levels impart a BAT proteome with greater capacity for oxidative metabolism and remodeling in response to early life exercise.

Prior research has shown inborn fitness dictates whole body and tissue specific metabolism, with LCR exhibiting lower VO_2max_, glucose tolerance, fat oxidation, and mitochondrial respiratory capacity, as well as increased serum triacylglyceride and insulin, greater blood pressure, and susceptibility to diet-induced obesity and metabolic diseases^23,24,26,49–53^. We have previously reported that adult HCR rats have greater glucose tolerance, lower body mass, and increased exercise capacity compared to LCR rats^26^. Further, we have also shown that HCR weanlings are smaller and leaner than LCR weanlings, while exhibiting clear differences in whole body and tissue level energeticssa^26,52^. Here, we further demonstrate that in addition to increasing TEE, early life exercise training increases BEE in both LCR and HCR rats when controlling for body mass. In contrast, strain alone did not appear to impact either BEE or TEE in sedentary rats. These data suggest that irrespective of inborn cardiorespiratory fitness, early life exercise increase both basal and total energy expenditure.

In the current study, we found that the mass of BAT depots were comparable in sedentary HCR and LCR rats, but when normalized to body mass HCR had larger relative BAT pads, supporting the purported link between BAT and increased cardiometabolic health. When we assessed BAT mitochondrial function through high-resolution respirometry, HCR-CTRL had greater UCP1-dependent respiration compared to LCR-CTRL, indicating greater levels of inborn fitness confer an increased BAT thermogenic capacity, suggesting that parental health and in particular parental CRF may program offspring BAT energetics.

Exercise also increased UCP1-dependent respiration in LCR, while exercise resulted in lower IC50 levels in both HCR-VWR and LCR-VWR compared to CTRL. Interestingly, UCP1 protein expression was not significantly altered by inborn fitness or VWR when examined through proteomics. Since thermogenesis in rodents is often associated with acute cold stress and/or exercise, a lack of change in protein abundance was not unexpected, though its pairing with increased functional capacity was interesting. Emphasizing exercise has significant therapeutic potential are our IC50 curve findings, where VWR led to significantly lower half-maximal inhibitory concentrations of GDP. In requiring less GDP binding to inhibit UCP1-driven respiration, our data suggest VWR may modulate UCP1 function *in vivo*. Indeed, UCP1 function *in vivo* is controlled by intracellular purine nucleotide (inhibition) and fatty acid (activation) levels, suggesting that UCP1 may be more responsive to acute changes in fatty acids and/or purine nucleotide levels. Additionally, VWR groups did not demonstrate the strain effect identified in CTRL rodents, further emphasizing the modulatory role of VWR in altered inhibitory purine nucleotide sensitivity. These results indicate BAT activation and functional capacity, rather than sheer abundance, may be more critical in overall metabolic health. Additionally, this finding underscores the necessity to directly examine mitochondrial function through respirometry, rather than simply using protein expression markers as proxies.

While cold exposure has been shown to activate BAT and UCP1 and increase BEE through non-shivering thermogenesis ^60–63^, prior research has shown exercise alone is able to alter adipose tissue morphology, vascularization, and thermogenesis^64–66^. While 10 weeks of high-intensity interval training exercise has been shown to increase BAT UCP1 levels in Wistar rats^67^, other groups have found treadmill or voluntary wheel running for up to six weeks failed to altered UCP1 levels in Sprague Dawley rats^62^ or obese mice^68^. Obesity alone has been shown to reduce BAT function, where Fu et al., attributed decreased VEGF in and increased adipose tissue hypoxia as the cause for impaired BAT function^68^. Our study found six weeks of voluntary wheel running exercise altered UCP1 capacity and function. Exercise resulted in smaller absolute BAT pad mass, regardless of strain, likely due to lower lipid content of BAT pads. Critically, early life exercise training resulted in greater UCP1 driven respiration and a lower GDP IC50 irrespective of inborn fitness, suggesting both greater thermogenic capacity of BAT mitochondria in response to early life exercise in addition to altered sensitivity to purine nucleotides, which play a critical role in regulating UCP1 function *in vivo*. Intriguingly, exercise mediated alterations in BAT UCP1 function occurred in concert with exercise mediated increases in both TEE and BEE. While greater TEE in response to exercise is likely a function of running volume, this is unlikely to be the case for BEE. While exercise may increase tissue remodeling and thus ATP demand, offering an explanation for greater BEE in rats that underwent early life exercise, altered UCP1 function and chronic sympathetic innervation with early life exercise may also explain some of the observed exercise effect on BEE.

Upon examination of the sedentary (CTRL) BAT mitochondrial proteome, HCR exhibited enrichment of fatty acid metabolism and catabolism processes when compared to LCR. Enriched BCAA catabolism, seen in our proteomics findings herein, has been shown to prevent glucose intolerance, limit diet-induced obesity, and increase thermogenesis^46,54^. Acad10, Slc25a1, Slc25a10, Acadsb, and Acad11, all mitochondrial proteins critical in amino acid metabolism, catabolism, and β-oxidation ^46,55–57^, were more abundant in HCR. Acad11 has been previously linked to elite performance in racehorses^58^, further emphasizing the link between enhanced oxidative capacity and greater genetically encoded cardiometabolic health. Our findings indicate BAT proteome remodeling in response to early life exercise are linked to inborn fitness levels. We found differences remained in protein expression pertaining to fatty acid metabolism between HCR-VWR and LCR-VWR. Further, HCR had a greater response to exercise when both exercised strains were compared to their sedentary controls. Recent findings indicate free fatty acids drive increased uncoupled respiration in white adipose tissue^69^. Increased fatty acid catabolism in HCR as compared to LCR may partially explain the remaining differences in cardiorespiratory fitness seen across strains. Increased substrate availability in HCR as opposed to increased thermogenic capacity could be key in explaining why exercise is only partially able to rescue measures of poor inborn fitness.

Unique to HCR-VWR was the abundance of DAPs pertaining to skeletal muscle. Both muscle progenitor cells and brown adipocytes come from Pax7^+^/Myf5^+^ progenitor cells^70^ and are involved in non-shivering thermogenesis^71^. Exercise has been shown to promote adipose tissue and skeletal muscle crosstalk, with myokine release promoting BAT thermogenesis^72^. With an exaggerated skeletal muscle related BAT proteome seen in HCR-VWR, it is possible increased crosstalk is driving elevated metabolic flexibility, lending itself to greater exercise tolerance and an increased response in terms of body composition. It is possible increased inborn fitness levels alone promotes increased crosstalk between the two tissues, suggested by increased Serhl2 expression in HCR-CTRL versus LCR-CTRL. Increased Serhl2 expression in exercised rat skeletal muscle is associated with increased lipid oxidation and triacylglycerol synthesis^73^, suggesting higher levels of inborn fitness confer greater lipid metabolism handling abilities. However, further research is necessary to confirm crosstalk between BAT and skeletal muscle as a potential mechanism by which increased inborn fitness levels exert phenotypic differences.

A limitation of our studies is that experiments were carried out at standard vivarium temperatures (24°C), while the rat thermoneutral zone is estimated to range between 28-31°C depending on strain^74^. However, this should not result in changes in strain- and exercise-effects seen across groups. Additionally, the offspring arm of this study was not powered to identify sexual dimorphisms, but rather strain- and exercise-effects. As such, these comparisons were not calculated, meaning it is possible there are sexual dimorphisms in the effect of exercise on modifying poor inborn fitness levels. Future directions include identifying the responsible parent in offspring metabolic health by cross breeding HCR and LCR strains. While this study examined the role of offspring exercise in metabolic health, additional studies will examine the role of parental exercise. Together, these studies will provide crucial information on the role of inborn fitness in lifestyle-modifiable cardiometabolic health.

## Summary

We report early-life exercise has the ability to partially rescue the deleterious metabolic phenotype imparted by low parental cardiorespiratory fitness. Further, six weeks of voluntary wheel running in early life increased BAT mitochondrial thermogenic capacity and function to a comparable degree. However, early life exercise resulted in greater enrichment of proteins associated with skeletal muscle hypertrophy and keto acid metabolism in HCR rats. These results, combined with our prior findings in liver and skeletal muscle, advocate for the consideration of inborn fitness in exercise training program implementation and optimization.

## Additional Information

### Data availability statement

All data is available from the corresponding author upon request

### Author Contributions

C.P., E.B., D.G.S., S.B., and L.K. conceived the study and designed the experiments; D.G.S., M.B., L.D.T., T.R., J.D.S., and C.P. carried out experimentation; D.G.S. carried out metabolic phenotyping and exercise interventions. M.S.K. analyzed and interpreted the rest of the data and wrote the manuscript. All authors reviewed, edited, and approved the final version of the manuscript.

## Acknowledgements

We thank the technical support of Mr. Trae Pittman and Mr. Bobby Fay of the Arkansas Children’s Nutrition Center’s animal vivarium. We also thank Samantha J. McKee at The University of Toledo for care and maintenance of the LCR/HCR rat colony and assistance in selecting breeding pairs for this study.

## Supplementary Information

Supplemental information can be found online in the Supporting Information section.

---

**Supplemental Figure 1.**
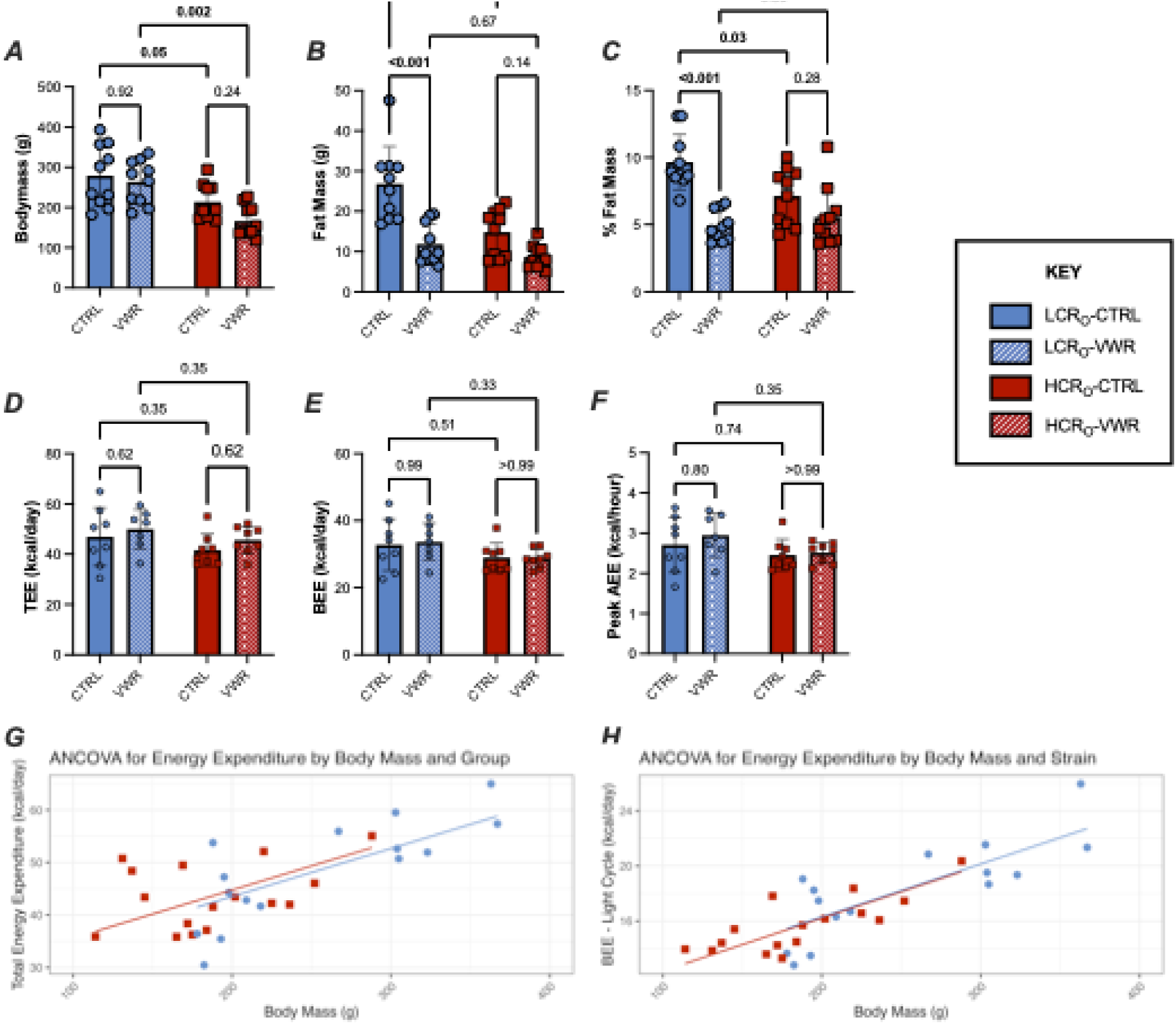
Inborn fitness and voluntary exercise alter physiology. (**A**) Body mass, (**B**) fat mass, and (**C**) relative fat mass at the time of euthanasia are shown. Average daily (**D**) TEE and (**E**) BEE are shown in addition to (**F**) peak AEE. ANCOVAs to compare (**G**) TEE and (**H**) BEE by body mass and strain are shown. Abbreviations: TEE—Total energy expenditure; BEE—Basal energy expenditure; AEE—Active energy expenditure; ANCOVA—Analysis of Covariance. Comparisons for A-F were made using two-way ANOVAs with multiple comparisons for main VWR and strain effects, where *P* < 0.05 was considered significant.

**Supplemental Figure 2.**
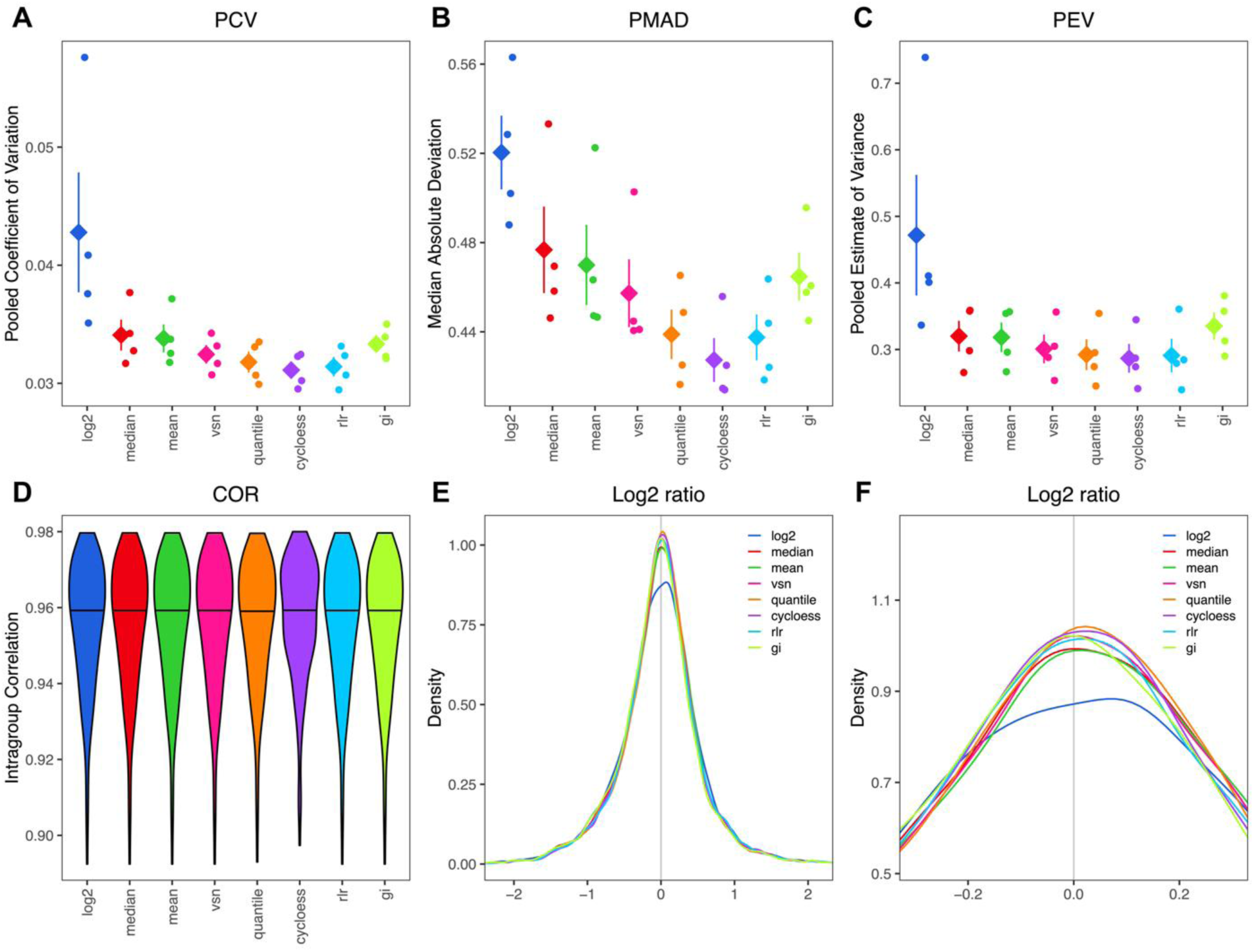
Normalization methods indicate cyclic loess is appropriate. Estimates of variability using PCV (a), PMAD (b), and PEV (c) are shown for the following normalization methods: log2, median, mean, vsn, quantile, cycloess, rlr, and gi. Intragroup correlation (d), in addition to density curves (e-f) are shown for each normalization method. Abbreviations: PVC: Pooled coefficient of variation; PMAD: Pooled mean absolute deviation; PEV: pooled estimate of variance; COR: correlation; VSN: variance stabilizing normalization; Cycloess: cyclic loess; RLR: robust linear regression; geospatial/general information.

